# Proinflammatory innate cytokines and metabolomic signatures shape the T cell response in active COVID-19

**DOI:** 10.1101/2022.03.11.483930

**Authors:** Akshay Binayke, Aymaan Zaheer, Jyotsna Dandotiya, Sonu K Gupta, Shailendra Mani, Manas Tripathi, Upasna Madan, Tripti Shrivastava, Yashwant Kumar, Anil K Pandey, Deepak K Rathore, Amit Awasthi

**Author notes:** Equal Contribution (Aymaan Zaheer, Jyotsna Dandotiya).

## Abstract

The underlying factors contributing to the evolution of SARS-CoV-2-specific T cell responses during COVID-19 infection remain unidentified. To address this, we characterized innate and adaptive immune responses with metabolomic profiling longitudinally at three different time points (0-3, 7-9, and 14-16 days post-COVID-19 positivity) from young mildly symptomatic active COVID-19 patients infected during the first wave in mid-2020. We observed that anti-RBD IgG and viral neutralization are significantly reduced against the Delta variant compared to the ancestral strain. In contrast, compared to the ancestral strain, T cell responses remain preserved against the delta and omicron variants. We determined innate immune responses during the early stage of active infection in response to TLR 3/7/8 mediated activation in PBMCs and serum metabolomic profiling. Correlation analysis indicated PBMCs-derived proinflammatory cytokines, IL-18, IL-1β, and IL-23, and the abundance of plasma metabolites involved in arginine biosynthesis were predictive of a robust SARS-CoV-2-specific Th1 response at a later stage (two weeks after PCR positivity). These observations may contribute to designing effective vaccines and adjuvants that promote innate immune responses and metabolites to induce long-lasting anti-SARS-CoV-2 specific T cells response.

## INTRODUCTION

T cell immune responses are indispensable in long-lasting protection against the coronavirus disease-19 (COVID-19)^1-3^. Although variants of concern (VoCs) acquired multiple mutations to evade the humoral immunity generated through vaccination, natural infection or both, T cell immunity remained largely preserved against variants of SARS-CoV-2^3-6^. Importantly, memory T cell response against SARS-CoV-2 persists for a long time since antigen exposure and thus protects from severe infection upon re-exposure to SARS-CoV-2 and its variants^7^. The frequency and magnitude of the antigen-specific T cell response could potentially affect the clinical manifestations of COVID-19^8-10^. Therefore, it is critical to understand the factors that shape the magnitude of T cell responses during SARS-CoV-2 infections.

Antigen-presenting cells (APCs) provide secondary signals in the form of cytokines that are critical in modulating T cell activation and differentiation^11^. However, the phenotype of cytokines secreted by innate immune cells in generating a comprehensive T cell response is not clearly understood in COVID-19. Systems biology studies have shown that proinflammatory cytokines, IL-6, IL-1β, and TNF-α, originating from the SARS-CoV-2 infected lungs but not from the peripheral blood cells, contribute to COVID-19 severity^12^. Therefore, studying the systemic levels of cytokines to predict the T cell response may prove misleading. There are limited reports of cellular innate immune responses and their correlation with adaptive immunity. Few recent longitudinal studies have documented the early innate and adaptive immune responses during mild COVID-19^13, 14^; however, no reports identify a correlation of an early innate immune response with the generation of T cell responses in active COVID-19. To address this, we used TLR 3, 7, and 8 agonists to stimulate PBMCs isolated within three days of PCR diagnosis of mild-COVID-19 patients and measured the range and magnitude of innate cytokine release, which were further correlated with the degree of virus-specific T cell response generated during active COVID-19 in mild cases where the virus was successfully cleared. We aim to shed light on the coordination of the innate and adaptive immune responses over the course of acute COVID-19 and understand the phenotypes of the immune cells involved at each stage to gain a holistic understanding of how different branches of the immune system act sequentially during infection.

We and others have reported that COVID-19 patients have altered metabolic pathways and dysregulation of energy production^15-19^. Since plasma metabolites substantially influence the immune cells in shaping protective immune responses, we tested whether levels of specific metabolites or metabolic pathways correlate with the antiviral immune responses. Our results reveal that the TLR-specific proinflammatory cytokine activity by PBMCs and a plasma metabolic signature correlate with a robust T-cell immune response against COVID-19 infection.

## MATERIALS AND METHODS

### Study Plan

To investigate the factors that contribute to the priming of a robust anti-SARS-CoV-2 T cell response, we longitudinally studied immune-metabolomic signatures during acute infection in young mildly symptomatic COVID-19 patients infected with the SARS-CoV-2 virus in mid-2020 by collecting blood samples on day 0-3 (indicated as “V1”), day 7 (indicated as “V2”) and day 14 (indicated as “V3”) from the date of PCR positive infection report. Clinically relevant medical information (e.g., age, patient-reported symptoms) was collected at the time of enrolment. For comparison, blood was also collected from age-matched SARS-CoV-2 RT-PCR-negative healthy volunteers (**Table S1**). The clinical cohort was considerably young and consisted of 8 females and 13 males with a median age of 28 years (IQR: 25:34) (**Table S1**). From the collected blood samples, we separated the plasma and PBMCs, which were cryopreserved until further experimentation. The humoral immune response against SARS-COV-2 and its variants was evaluated using the plasma samples collected from the 21 patients. PBMCs from all three-time points were used to determine antigen-specific T cell response dynamics during acute COVID-19. To study the status of innate immunity against SARS-COV-2, we *ex vivo* stimulated the PBMCs of V1 samples with a cocktail of TLR 3/7/8 agonists, and the released cytokines were quantified. We performed a Spearman correlation analysis of the innate immune response from V1 samples with the SARS-CoV-2 specific T cell immune response of V3. The metabolic signature obtained from the plasma was further correlated with the SARS-CoV-2 specific innate and T cell immune responses (**Fig. 1A**).

**Fig. 1:**
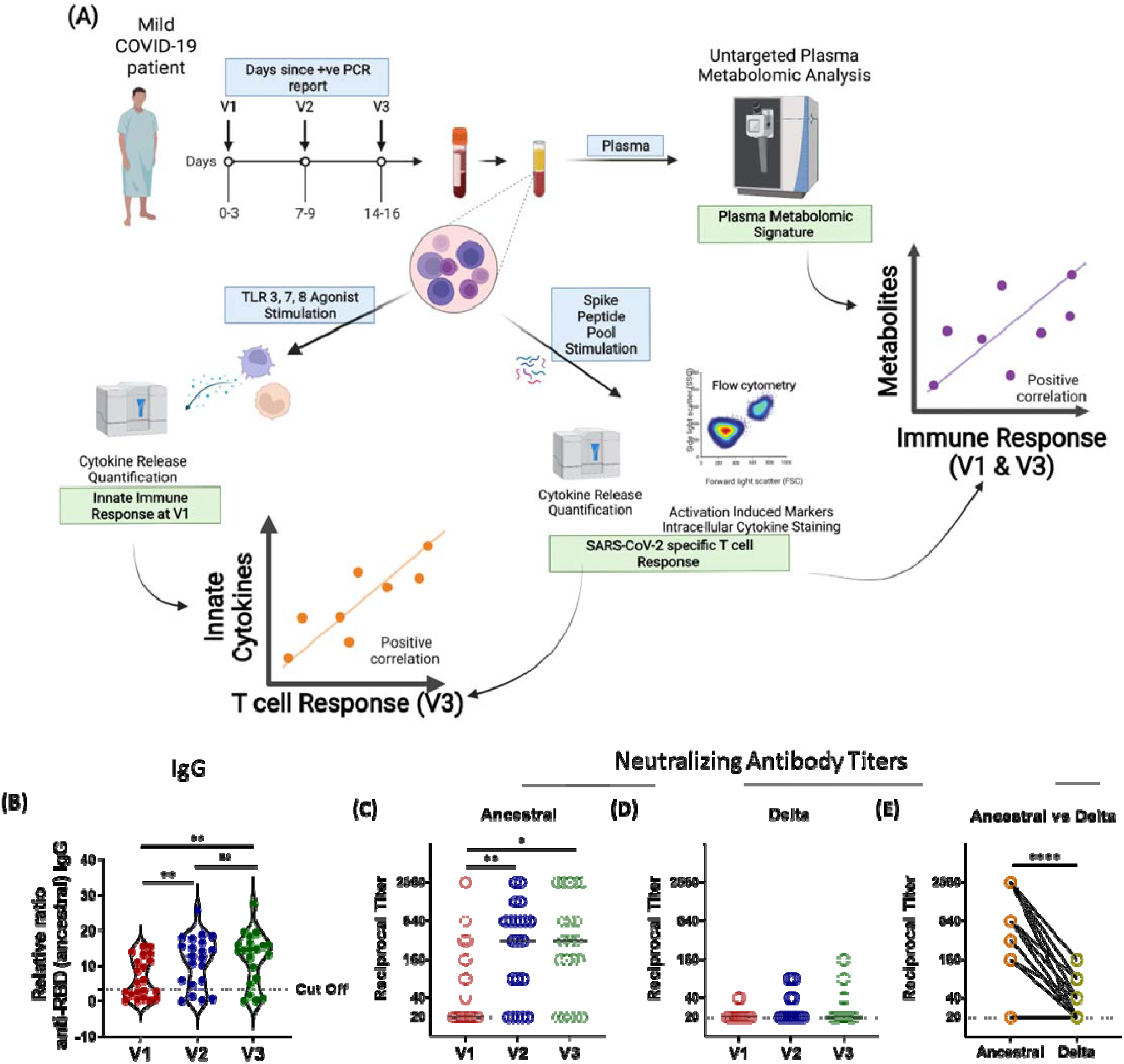
Longitudinal Analysis of Humoral Immune Response against the SARS-CoV-2 during acute COVID-19 infection. **(A)** Graphical representation of the study plan **(B)** The longitudinal anti-RBD IgG responses were evaluated by performing ELISA against the RBD proteins of the Ancestral (Wuhan isolate). All data, represented as ratio-converted ELISA reads to a pool of pre-pandemic negative control samples (relative ratio), were plotted using violin plots. **(C-E)** The longitudinal neutralizing antibody titers against **(F)** the ancestral strain and **(G)** the delta strain of SARS-CoV-2 during V1 (day 0-3), V2 (day 7), and V3 (day 14) from COVID-19 positivity **(H)** The paired representation of NAb titers in the active COVID-19 patients during V3 against the ancestral and delta variants of SARS-CoV-2. The dots represent each individual sample. Tukey’s multiple comparisons and two-sided Wilcoxon Signed Rank tests were employed for unpaired and paired analysis, respectively. n.s = not significant, *= p ≤ 0.05, **= p ≤ 0.01, **** = p < 0.0001.

### Human Ethics

All the experiments were performed according to the suggested guidelines of the Institutional Ethics Committee (Human Research) of THSTI and ESIC Hospital, Faridabad (Letter Ref No: THS 1.8.1 (97) dated 07th July 2020). Peripheral blood samples were collected from asymptomatic or mildly symptomatic COVID-19-positive and healthy individuals after receiving written informed consent. Individuals were enrolled in this study upon appropriate approval from the Institutional Ethics Committee (Human Research) of THSTI.

### THSTI In-house RBD IgG ELISA

ELISAs to detect IgG binding to receptor binding domain (RBD) were performed as previously described^20^. Positive convalescent and negative control samples were added to each plate for normalization. The assay results were normalized by dividing the blank subtracted readings of each sample by negative control to obtain the fold change reading.

### Virus Neutralization Assay

Virus microneutralization assay titers were estimated as described previously ^12^. Briefly, plasma samples were serially diluted from 1/20 to 1/2560 and incubated with the ancestral (Wuhan isolate) and the delta (B.1.617.2) SARS-CoV-2 isolates. 50% neutralization values were estimated with four-parameter logistic regression.

### Peptide Pool

15mers peptide pools with an overlap of 11 amino acids spanning the entire sequence of Spike protein (157+158 peptides, JPT, PepMix, Cat. No.PM-WCPV-S-1 (Ancestral), PM-SARS2-SMUT06-1 (Delta) and PM-SARS2-SMUT08-1 (Omicron)) were used for the determining the ancestral, B.1.617.2 and B.1.1.529 SARS-CoV-2-spike specific T-cell responses by AIM assay, ICS and cytokine bead assay.

### SARS-CoV-2 Specific T cell response

SARS-CoV-2 Specific T cell responses were studied as described previously^6, 21, 22^. PBMCs were stimulated with peptide pools at a 2 μg/ml/peptide concentration and with an equimolar dimethyl sulfoxide (DMSO) concentration as a negative control. Phytohemagglutinin (PHA, Roche; 5μg/ml) was used as a positive control. As a co-stimulant, anti-CD28 and anti-CD49d were added (BD Biosciences, USA). The cells were cultured for 22-24h, and monesin (GolgiStopTM, BD Bioscience, USA) was added during the last six hours. After stimulation with peptides, the culture supernatant was separated and stored at <-20°C until further use, and the cells were stained for flow cytometry. Samples were acquired on a fluorescence-activated cell sorter (FACS) Symphony™ instrument (BD Biosciences) using BD FACSuite software version 1.0.6 and analyzed with FlowJo software version VX (FlowJo LLC, BD Biosciences). Functional profiles were deconvoluted by employing Boolean gating in FlowJo version XV. Spike-specific cytokine production was background-subtracted by the values obtained with the DMSO-containing medium, and the negative values were set to the Limit Of Detection (LOD). The stimulation index (SI) and LOD were calculated as described previously ^22^; briefly, the SI was calculated by dividing the frequency of AIM+ CD4 or CD8 cells recorded in stimulated wells by the unstimulated wells.

### Metabolomic Analysis

Metabolomic profiling of plasma metabolites was performed as described previously^23-26^. The LC/MS obtained data were processed using the Progenesis QI for metabolomics (Nonlinear dynamics, a Waters Company) software using the default setting. Manual processing of the raw data by removing drugs and exposome metabolites led to the identification of 176 metabolites at all three-time points. All the statistical and functional analysis, including the PCA, heat map, pathway enrichment analysis, PLS-DA, and analysis of variance (ANOVA), was done based on the observed peaks intensity using the online open-source software Metaboanalyst 5.0. Before analysis, a data integrity check was performed, and the raw data were normalized by sum, log-transformed, and scaled by Pareto scaling. The volcano plots were constructed using the online available package VolcaNoseR ^27^.

### In vitro stimulation of PBMCs with TLR agonists

PBMCs were stimulated as previously described ^12^, with some modifications. Approximately one million cells were added to each well, and all wells received stimulation with a viral cocktail containing 4.0 μg/ml R848 and 25 μg/ml poly I:C. Negative control DMSO stimulated cells were also cultured for each sample. The cells were incubated in a 5% CO2 incubator at 37ºC for about 24 hours, after which the culture supernatant was collected to analyze cytokine secretion.

### Analysis of Cytokine Secretion by Luminex

Cytokine secretion in the cell culture supernatant was analyzed with a customized Premixed Multi-Analyte Luminex Discovery Assay kit (LXSAHM-10 (peptide pools stimulation); LXSAHM-11 (TLR agonist stimulation), R&D systems). The assays were performed as per the manufacturer’s instructions. The analytes examined upon peptide pool stimulation were Granzyme B, IL-2, IL-5, IL-12 p70, IL-23, IFN-gamma, IL-4, IL-6, IL-17/IL-17A, and TNF-alpha, whereas the analytes examined upon TLR stimulation were IL-23, IFN-gamma, IL-6, IL-12 p70, TNF-alpha, IL-33, IL-8, IL-10, CCL2, IL1-beta, and IL-18.

### Data Representation and Statistical Analysis

The statistical analysis and data representation was performed using GraphPad Prism 9.0 and FlowJo XV unless otherwise stated. The antibody responses were compared using the RM one-way ANOVA Tukey’s Multiple Comparison Test. The cytokine and T cell response against each variant versus ancestral strain and at different stages of acute infection was calculated and compared using the Wilcoxon matched-pairs signed-rank test. T cell responses were calculated as background-subtracted data by subtracting the values obtained from the SARS-CoV-2 peptide pool stimulation from the DMSO stimulation. Negative values were set to the LOD. The cytokine secretion response upon peptide or TLR stimulation was calculated as fold change data for the correlation analysis by dividing the stimulated wells by unstimulated wells of the same sample. The correlation between the innate and adaptive immune response was performed using the Spearman correlation test. For metabolomic analysis, processing the raw data by removing drugs and exposome metabolites led to the identification of 176 metabolites at all three stages (Visit 1, Visit 3, and Healthy Control). All the statistical and functional analysis, including the PCA, heat map, pathway enrichment analysis, PLS-DA, and analysis of variance (ANOVA), was done based on the observed peaks intensity using the online open-source software Metaboanalyst 5.0. Before analysis, a data integrity check was performed, and the raw data were normalized by sum, log-transformed, and scaled by Pareto scaling. The volcano plots were constructed using the online available package VolcaNoseR ^27^.

A broadscale Spearman correlation analysis of the metabolites belonging to the V1 and V3 groups with the innate and adaptive immune responses of V1 and V3 was performed. Those metabolites belonging to V1 and V3 that significantly correlated (p<0.05, r>0.45) with the proinflammatory innate immune responses and/or TH1 skewed T cell immune responses and has pathway library of KEGG were selected for Pathway Analysis using the Metaboanalyst 5.0 software ^28^. The graphical abstract and figure 1A were prepared using Biorender.com.

## RESULTS

### Humoral Immune responses against SARS-CoV-2 variants is broadly inadequate in active-COVID-19 patients

To study the kinetics of the antibody response against the SARS-CoV-2, we first investigated the IgG response against the SARS-CoV-2 RBD protein using Enzyme-linked immunosorbent assays (ELISAs). For all assays, blank-subtracted colorimetric values were normalized to a pre-pandemic negative control plasma sample added to each assay plate and expressed as ratios to this pool of negative samples as described previously^29^. We observed that the anti-RBD (ancestral) IgG titers increased about three-fold by V2 and V3 compared to V1 (**Fig. 1B**). However, similar to the previous reports in vaccinated and convalescent individuals^30^, the cross-reactive antibody response against the RBD proteins of the delta (B.1.617.2), beta (B.1.315), and alpha variants (B.1.1.7) was significantly reduced (**Fig. S3**). While the median neutralizing antibody (Nab) titers increased significantly by V2 and V3 as compared to V1 against the ancestral virus (**Fig. 1C**), cross-reactive NAb titers against the delta variants were found to be significantly decreased at V2 and V3 as compared to the ancestral virus (p<0.0001, **Fig. 1E**). We observed that even by V3, only 33% (7/21) individuals had detectable levels of cross-reactive Nabs against the Delta variant (**Fig. 1D**). Therefore, in concurrence with the previous reports, the cross-reactive antibody-mediated protection against different variants of SARS-CoV-2 in active COVID-19 patients infected with ancestral strain is broadly inadequate.

### Spike-specific T cell response in COVID-19 patients

We studied the antigen-specific T cell response from the PBMCs of active COVID-19 patients by stimulating them with peptide pools spanning the entire length of the spike protein for 20-24h. The T cell response upon peptide pool stimulation was measured by calculating the expression of T cell receptor (TCR)-dependent activation-induced markers (AIM) ^10^ and Th1 cytokines after background subtraction from DMSO stimulated wells ^31^. The CD4+ T cells co-expressing CD137 and OX-40 are designated as CD4+AIM+ cells (**Fig. 2A**), and the CD8+ T cells co-expressing CD137 and CD69 are defined as CD8+AIM+ cells (**Fig. 2B**) ^10, 21^.

**Fig. 2.**
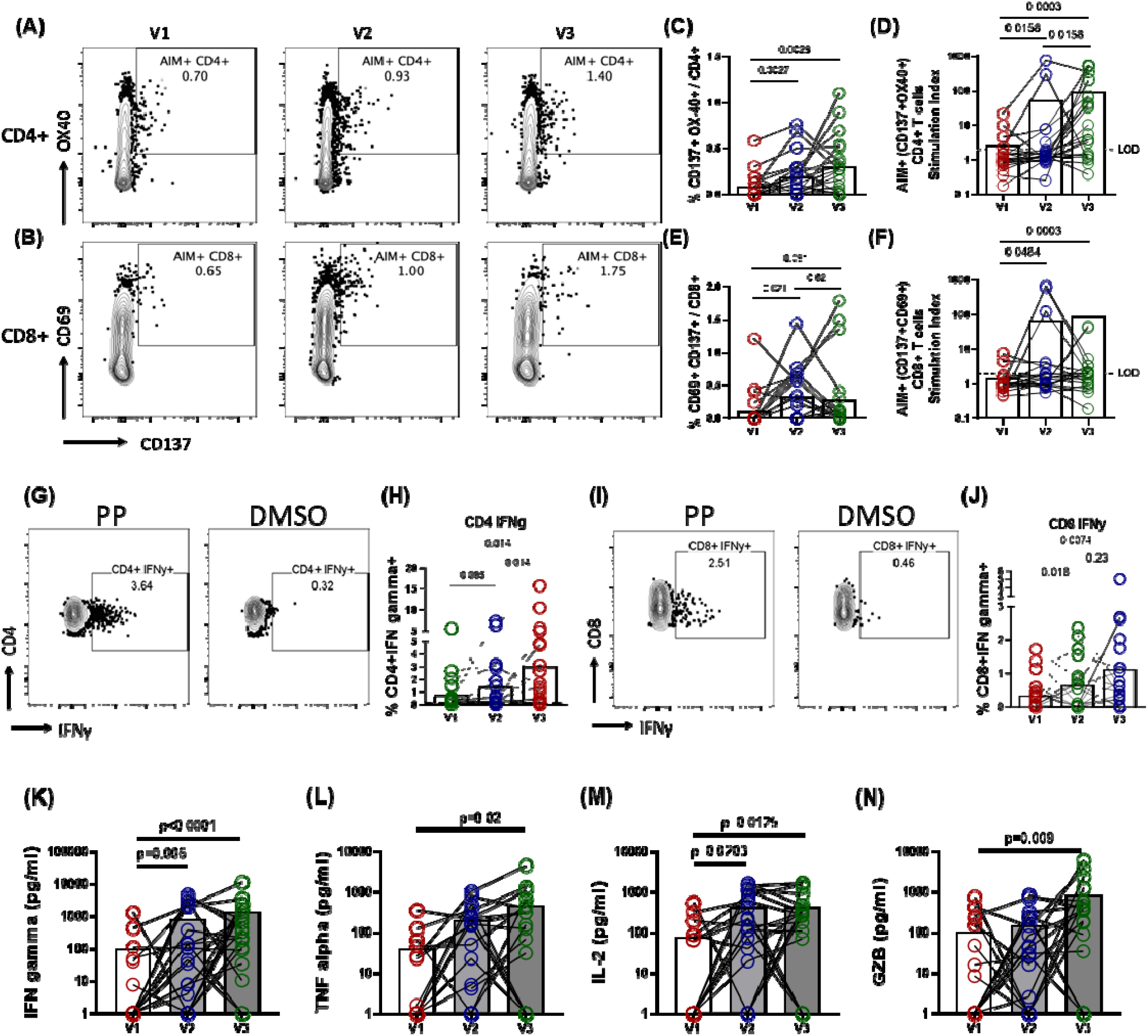
Longitudinal dynamics of antigen-specific T cell immune responses during COVID-19 infection. Representative flow cytometry plots of SARS-CoV-2 Spike specific T cells expressing activation-induced markers (AIM) **(A)** CD4+ (CD137+OX40+) **(B)** CD8+ (CD137+CD69+); **(C-F)** Longitudinal analysis of the AIM response in paired samples from the same subject **(C)** percentage frequency of CD4+AIM+ cells **(D)** Stimulation index (SI) of CD4+ AIM+ cells **(E)** %frequency of CD8+AIM+ cells **(F)** SI of CD8+ AIM+ cells. **(G, I)** Representative flow cytometry plots for the intracellular cytokine staining (ICS) assay of cells upon stimulation with spike peptide pool compared to the DMSO; Longitudinal analysis of the frequency (percentage of total CD4+ or CD8+ cells) of IFN-γ+ cells in paired samples from the same subject. **(H)** CD4+ Interferon gamma (IFNγ) **(J)** CD8+ Interferon gamma; Each datapoint shown is background subtracted (DMSO stimulated wells), and bars represent mean values for each time point. Two-sided Wilcoxon Signed Rank tests were employed for paired non-parametric analysis. LOD = Limit of detection for AIM assay was SI<2. **(K-N)** Longitudinal dynamics of antigen-specific cytokine release by PBMCs during COVID-19 infection **(K)** IFNγ **(L)** TNFα **(M)** IL-2 **(N)** Granzyme B. Graphs represent longitudinal cytokine released (background subtracted) by paired PBMC samples from the same subject upon stimulation with SARS-CoV-2 spike peptide pool for 22-24h. Bars represent mean values.

The activation-induced markers (AIM) in CD4+ T cells were observed as early as V1 (6/21), which may be attributed to pre-existing cross-reactive T cells from prior common cold coronavirus infections^32^. The magnitude of SARS-CoV-2-specific CD4+ T cells increased significantly by V2 (day 7, p = 0.0027) and by V3 (day 14, p = 0.0028) (**Fig. 2C**). Consistent with the previous reports^10^, the AIM+ cells in CD4+ T cells were observed in ∼76% of the patients by V3 (16/21) (**Fig. 2D**). Likewise, AIM+ spike-specific CD8+ T cells were detected as early as V1 (4/21). Interestingly, unlike CD4+ T cells, the frequency and magnitude of AIM+ cells in CD8+ T cells peaked quickly from V1 to V2 (p = 0.021) and did not change significantly from V2 to V3 (p = 0.52) (**Fig. 2E**). Antigen-specific CD8+ T cells were detected in only 57% (12/21) (**Fig. 2F**) of the patients at V3, in line with the previous findings reported by Moderbacher et al., 2020 ^10^.

The functional antigen-specific T cell response was tested by intracellular expression of IFN-γ and IL-2 by performing intracellular cytokine staining of spike peptide pool stimulated PBMCs. Similar to the CD4+ AIM+ response, the CD4+ IFN-γ+ cells frequency increased significantly from V1 to V3 (p=0.014, **Fig. 2G-H**). Likewise, the functional SARS-CoV-2 specific cytotoxic T cell frequency characterized by CD8+ IFN-γ+ cell frequency increased significantly from V1 to V3 (p=0.0074, **Fig. 2 I, J**). Therefore, to summarize, we found a steady increase in the CD4 and CD8 AIM and functional T cell response, which was identified by intracytoplasmic expression of IFN-γ in both CD4+ and CD8+ T cells.

To further understand the range of cytokines released after stimulation of PBMCs with peptide pool, the release of cytokines was calculated by cytokine bead assay. With the progression of time, we observed a steady increase in the secretion of Th1 specific cytokines such as IFN-γ (mean: V1:100 pg/ml; V2: 810 pg/ml; V3: 1352 pg/ml, **Fig. 2K**), TNFα (mean: V1: 40 pg/ml; V2: 206 pg/ml; V3: 478 pg/ml, **Fig. 2L**), and IL-2 (mean: V1: 77 pg/ml; V2: 430 pg/ml; V3: 429 pg/ml, **Fig. 2M**), whereas Th-2 and Th-17 specific cytokines did not change significantly (**Fig. S7 A, C, D)**. We did not detect any notable differences in the IL12p70 secretion **(Fig. S7 E)**, but as the antigen-specific T cell response increased, the proinflammatory cytokines IL-23 and IL-6 upon antigen exposure increased significantly by visit 3 (IL-6: mean:898 pg/ml; p=0.013; IL-23: mean: 711 pg/ml; p=0.034, **Fig. S7 B, F)**. The cytotoxic response, evaluated by the levels of Granzyme B, also increased extensively at V3 compared to V1 (p=0.009, mean V3= 841 pg/ml, **Fig. 2N**).

To sum up, the SARS-CoV-2 specific T cell immune responses were predominated by Th1 cytokine expression, and their magnitude progressively increased with the advent of time since antigen exposure.

### Cross-reactive spike-specific T cell response against delta and omicron variants is largely preserved in active COVID-19 patients

We found that the antigen-specific T cell response is significantly increased by 14 days post detection of COVID-19 infection with the ancestral strain. To confirm whether T cell responses elicited by the ancestral strain in active COVID-19 patients could cross-react with the delta and omicron spike protein, we tested and compared the T cell response in the PBMCs (n=24) of active COVID-19 patients exposed to the ancestral strain of SARS-CoV-2 from V2 and V3 against the spike peptide pools of the ancestral, delta, and omicron variants (**Fig. 3**). We observed that the cross-reactive T cell response is largely preserved with a slight reduction against delta (1.3-fold) and the omicron (1.5-fold) spike protein compared to the ancestral spike. Compared to the ancestral response, the geometric mean of the AIM+ in CD4+ response was reduced by 31% against delta (p=0.0162) and 58% against Omicron spike (p=0.0005) (**Fig. 3A**). However, the functional profile remained broadly comparable among the variants as determined by the expression of intracellular cytokines IFN-γ, IL-2, and TNFα (**Fig. 3B-D**).

**Fig. 3.**
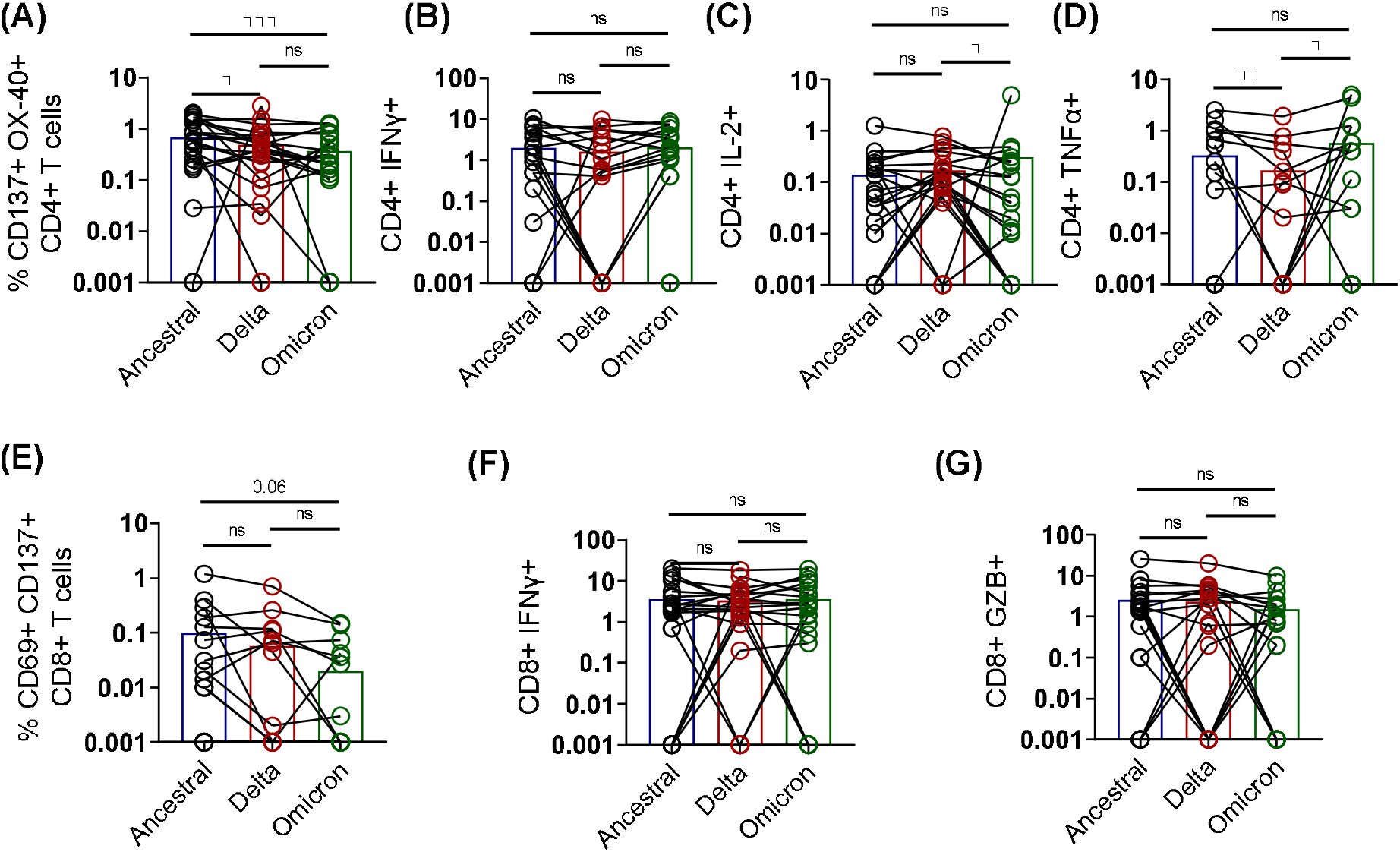
Cross-reactive CD4+ and CD8+ T cell response against Delta and Omicron Spike in acute. COVID-19 patients Spike-specific T cell responses in selected samples of active COVID-19 patients (n=24) were simultaneously tested for reactivity against spike peptide pools of Ancestral, Delta (B.1.617.2), and Omicron (B.1.1.529). **(A-D)** Frequencies of antigen-specific CD4+ T cell responses **(A)** CD4+ AIM [CD137+OX40+] **(B)** CD4+ IFNγ **(C)** CD4+ IL2 **(D)** CD4+ TNFα. **(E-G)** Frequencies of antigen-specific CD8+ T cell responses **(E)** CD8+ AIM [CD137+CD69+] **(F)** CD8+ IFNγ **(G)** CD8+Granzyme B Bars represent mean values, each dot represents an individual sample, and the solid line connects the same sample stimulated with different peptide pools. Two-sided Wilcoxon Signed Rank tests were employed for paired non-parametric analysis. AIM, activation-induced markers; GZB, granzyme B; ns, non-significant

Although the geometric mean of antigen-specific CD8+ AIM response was reduced by three and five-fold in the delta and omicron variants, respectively, the difference was non-significant (**Fig. 3E**). Moreover, the functional phenotypes of the antigen-specific CD8 T cells were comparable among the VoCs compared to the ancestral spike antigen (**Fig. 3 F, G**).

Taken together, in concordance with the previous reports^3-6^, the cross-reactive T cell immune responses persist despite the humoral immune response being abrogated against the VoCs in actively infected individuals. Our results imply the importance of the cellular immune responses against COVID-19. Thus it is essential to understand further what factors determine the generation of a robust T cell response during active COVID-19.

### Early proinflammatory innate cytokine response correlates with a robust T cell response

Next, we tested which innate immune responses correlate with the adaptive immunity, especially T cell immune response in active COVID-19 patients. Given the earlier findings that COVID-19 infection impairs the APCs such as the pDCs and myeloid cells^12, 33^, we wanted to test what functional phenotype of APCs correlates with the virus-specific T cell responses. We performed *ex vivo* stimulation of V1 PBMCs of our COVID-19 cohort with a cocktail of synthetic agonists of toll-like receptor (TLR) 3, 7, and 8, which are known to sense virus-derived molecules and initiate an antiviral response^34^. In the presence and absence of the TLR agonists cocktail of poly I:C (TLR3) and R848 (TLR 7/8), the PBMCs were cultured for 24h, as described previously^12^. The release of cytokines such as IL-18, IL-23, IL-1β, IL-6, IL-33, IL-8, IL-10, CCL2, IL12p70, and IFNγ, in the culture supernatant was estimated by a magnetic bead-based cytokine assay. The cytokines levels in TLR stimulated wells were significantly higher than in the unstimulated wells (**Fig. S2**). Therefore, to account for the differences in the number of PBMCs cultured, fold change in the cytokine response was evaluated by dividing the cytokine response in stimulated wells by unstimulated wells. The fold change in cytokine expression of TLR stimulated samples compared to the unstimulated wells was correlated with the antigen-specific T cell responses such as the AIM markers, ICC-cytokines, and the cytokines expressed in the culture supernatant (such as TNFα, IFNγ, IL-6, IL-4, IL-17, IL-2, IL-5, IL-12p70, GZB, and IL-23 by PBMC samples from V3 upon peptide pool stimulation. (**Fig. 4A**).

**Fig. 4.**
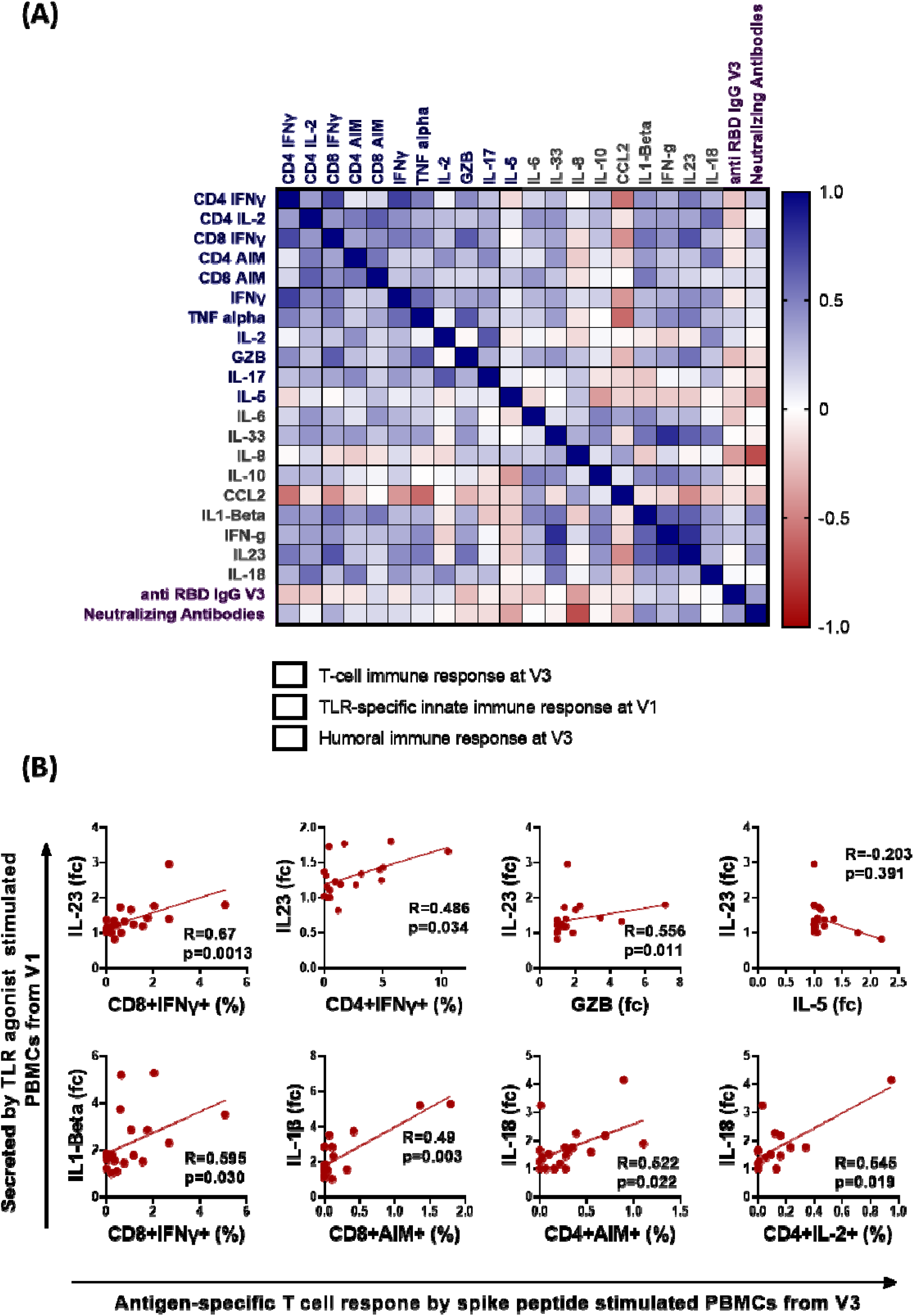
Correlation of early innate and adaptive immune responses in active COVID-19 patients. Correlation between the cytokine released upon TLR agonist stimulation of PBMCs from early stages of COVID-19 infection (V1) and the magnitude of adaptive immune responses during the later stages of COVID-19 disease (V3). **(A)** A heatmap of spearman correlation analysis; the blue rectangle depicts the analysis of stimulated PBMCs using flow cytometry or the fold change of cytokines released in the culture supernatant upon peptide stimulation; the grey rectangles represent the fold change of cytokines released in the culture supernatant upon stimulation with TLR agonists: poly I:C (25 μg/ml) and R848 (4 μg/ml) for 24h and the lavender color rectangle represents the humoral immune response at V3 **(B)** XY correlation plots where the y axis represents the fold change of cytokines secreted upon TLR agonist stimulation of PBMCs from V1, and the x-axis represents the antigen-specific T cell response evaluated upon peptide pool stimulation of PBMCs from V3. Spearman correlation coefficient (R) is indicated. fc, fold change – obtained by dividing the cytokine concentration in stimulated wells by unstimulated wells; GZB, granzyme B

The intracellular cytokine levels were highly correlated with the cytokines in the culture supernatant, indicating the robustness of the sample analysis (**Fig. 4A**). For example, the CD4+ and CD8+ T cell IFN-γ significantly correlated with the IFN-γ levels in the culture supernatant (CD4: r =0.751, p=0.00014; CD8: r=0.522, p=0.0181; **Fig. 4A**). The Spearman correlation analysis showed that the PBMCs from individuals that secreted a heightened proinflammatory cytokine response in the form of IL-18, IL-1β, and IL-23 upon TLR stimulation during the early time-point of infection (V1) correlated significantly with the spike-specific T cell immune response in the form of the expression of CD4 and CD8 IFN-γ, CD4 and CD8 AIM, and Granzyme B (GZB) secretion (**Fig. 4B**). In contrast, the remaining cytokines such as IL-8, CCL2, and IL-10 did not correlate. Moreover, the proinflammatory cytokines did not associate with non-Th1 cytokines such as IL-5, and IL-17 and humoral immune responses (**Fig. 4A, B**). Although we observed a significant negative correlation between the neutralizing antibody response and the fold change in IL-8 secretion upon TLR stimulation (r=-0.7, p=0.0012), the observations were not further evaluated because IL8 expression was not significantly altered upon TLR cocktail stimulation (**Fig. S2**). To sum up, individuals exhibiting a proinflammatory functional phenotype of innate immune cells during the early stages of infection tend to develop robust antiviral T cell immunity.

### Distinct Metabolomic Alterations characterize acute COVID-19 infection

The correlation between metabolic profile with innate and adaptive T cell response in active COVID-19 infection remains unexplored. Moreover, altered metabolites in active COVID-19 patients might help predict the disease outcome and anti-SARS-CoV-2 immune responses. To determine the metabolic landscape in active COVID-19 patients, we performed a plasma metabolomic analysis to identify changes throughout COVID-19 infection. Multiple reports have identified metabolomic alterations during COVID-19^35-38^. To obtain a broad view of the metabolic status, we studied the longitudinal plasma metabolite profile in our cohort of 21 COVID-19 patients and compared them with the plasma of 12 healthy volunteers (**Table S1**). The plasma samples from V1 and V3 were processed for UHPLC-MS/MS analysis, and 240 different metabolites were identified. These 240 metabolites were further screened to remove drugs and other artificial compounds and derivatives to obtain 176 metabolites. Unsupervised PCA analysis of the metabolites in our patient cohort displayed a clear distinction that separated the control (healthy individuals), COVID-19 early time point (V1), and late time point (V3) individuals (**Fig. 5A**). The Partial Least Squares – Discriminant Analysis (PLS-DA) was employed to identify the top-15 important metabolites across the three different groups (**Fig. 5B**). Moreover, hierarchical clustering of the top 50 metabolites showed a notable shift in the metabolic signature among acute COVID-19 patients compared to the healthy individuals (**Fig. 5 C**). The PLS-DA and hierarchical clustering analysis pointed out that the metabolites and derivatives of the Citric Acid cycle, such as Citrate, Itaconate, and trans-aconitate, were significantly reduced in COVID-19 patients (**Fig. 5B-C**). In contrast, metabolites such as glutamic acid derivatives – Pyroglutamate, and N-methyl-L-glutamic acid were enriched in the plasma of COVID-19 patients (**Fig. 5B-C**). Of the 176 metabolites, 158 were significantly differentiated in COVID-19 patients (FDR<0.001), exhibiting a distinct metabolic profile. Pathway enrichment analysis using the KEGG database showed the metabolites involved in the Phenylalanine metabolism, Arginine and Proline metabolism, etc., were significantly altered during COVID-19 (**Fig. 5D**). While comparing the metabolic profile during the early and later stages of infection, 33 metabolites were significantly down and 13 significantly up (p=0.001, FC>1.5, **Fig. 5E**).

**Fig. 5.**
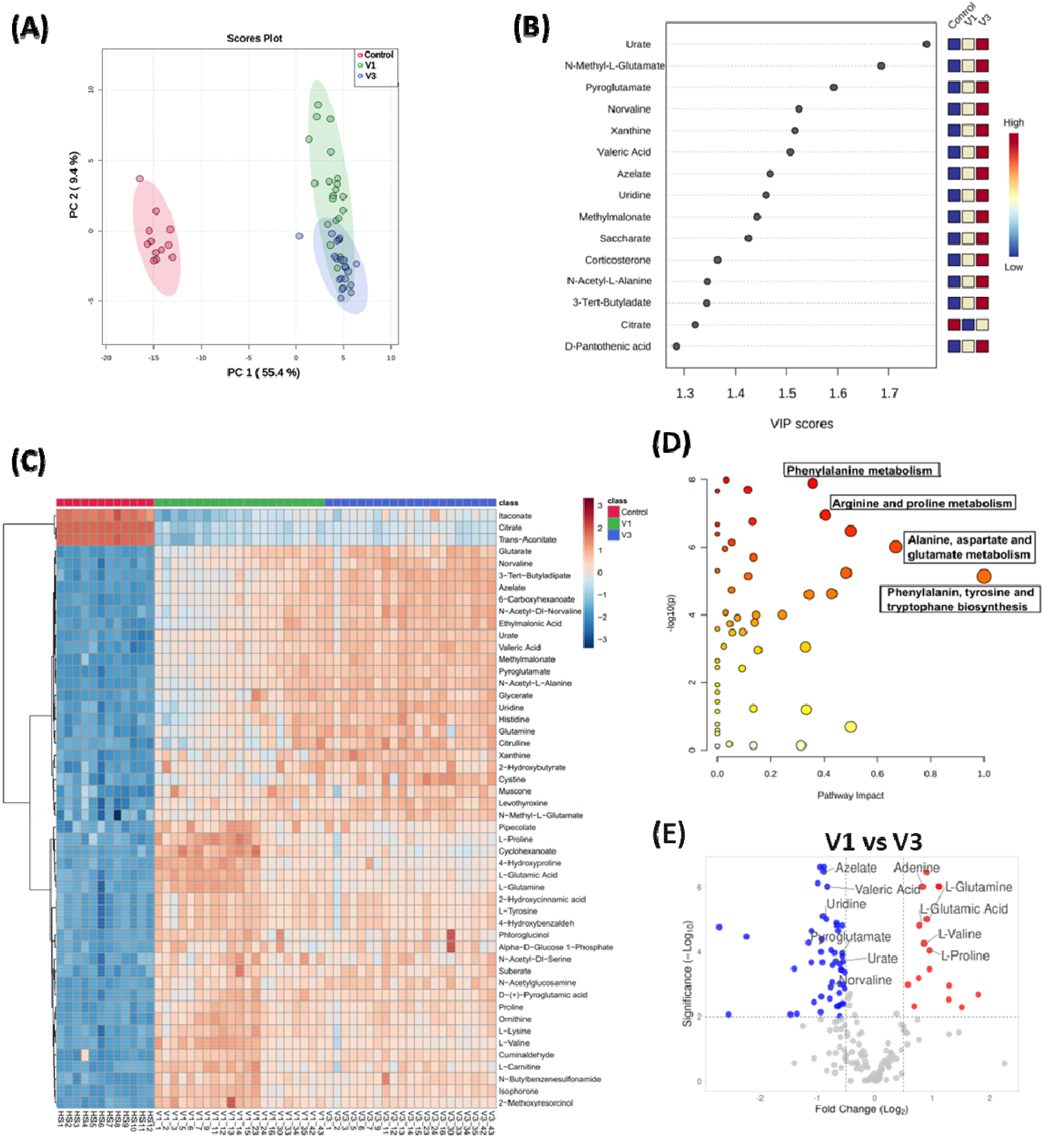
Metabolomic analysis of SARS-CoV-2 patient plasma samples compared to age-matched healthy controls. **(A)** Principal Component Analysis of the untargeted metabolites of the three groups **(B)** Important features identified by PLS-DA. The colored boxes indicate relative concentrations of designated metabolites in each study group. **(C)** hierarchical cluster analysis is shown as a heatmap of the top 50 metabolites (FDR<0.05). **(D)** Pathway enrichment analysis using KEGG of all the metabolites (n=158) significantly altered during COVID-19 infection. **(E)** Volcano plot comparing the significantly changed (p=0.001, FC>1.5) metabolites during the early (V1) and later stages of disease (V3).

To understand the immune-metabolomic interaction, we performed a Spearman correlation of metabolites with the innate and adaptive immune responses in COVID-19 patients and shortlisted the significantly correlating metabolites (r>0.45, p<0.05). Pathway analysis of plasma metabolites that significantly correlated with the virus-specific innate immune response from V1 showed enrichment of metabolites involved in arginine biosynthesis, D-glutamine, and D-glutamate metabolism, glutathione metabolism, histidine metabolism, etc. (**Fig. 6A**). Similarly, pathway analysis of metabolites that significantly correlate with the antigen-specific T cell responses at V3 displayed enrichment of metabolites involved in arginine biosynthesis, arginine and proline metabolism, purine metabolism, etc. (**Fig. 6B**). Notably, the normalized levels at V1 and V3 of metabolites involved in arginine and proline biosynthesis and metabolism, such as L-arginine, L-proline, L-ornithine, L-citrulline, α-ketoglutarate, L-glutamine and L-glutamate were significantly correlated with the innate and T-cell immune responses (**Fig. 6C**).

**Fig. 6.**
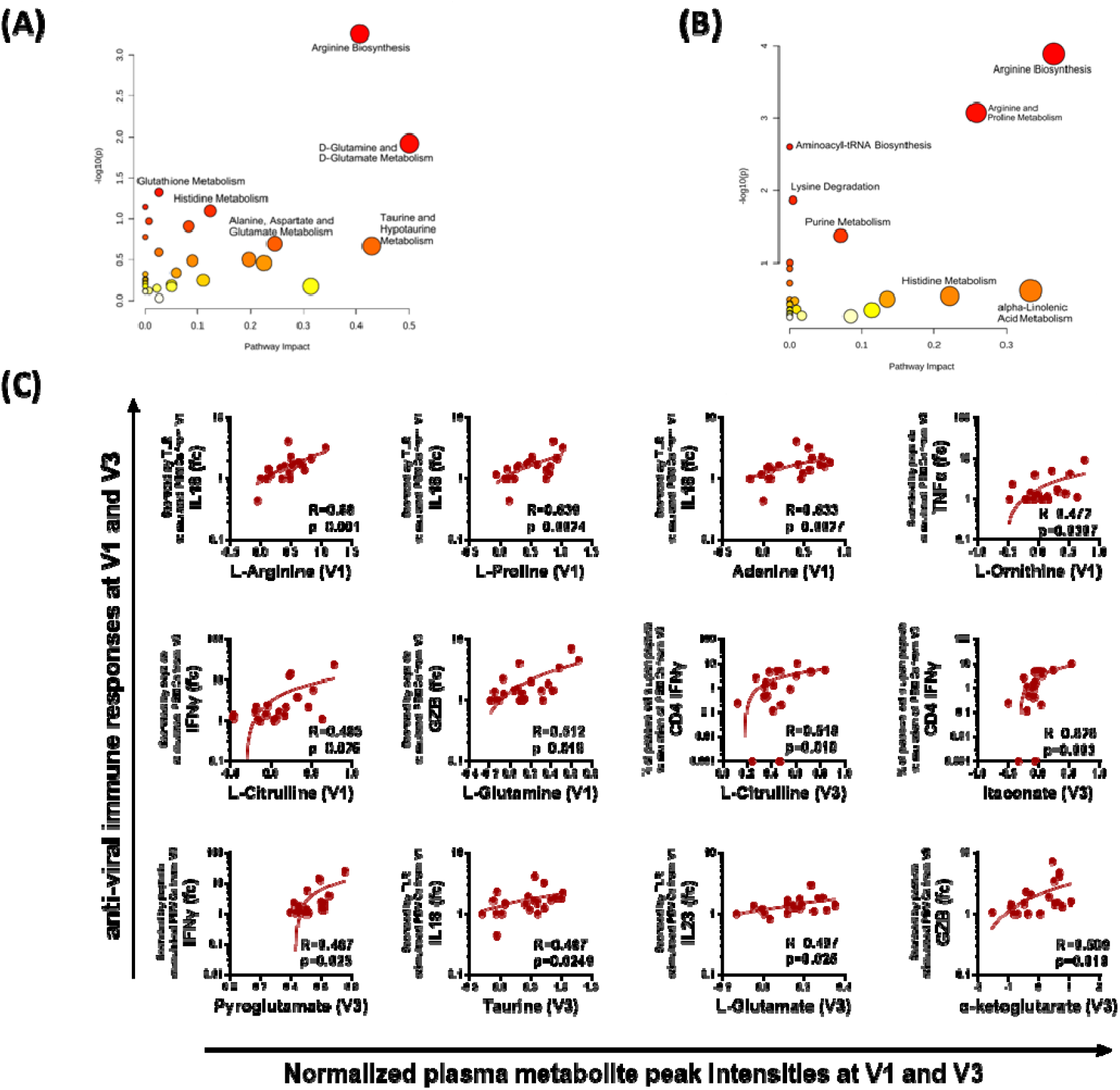
Correlation of metabolites with immune responses. Pathway analysis of plasma metabolites that significantly correlated (r>0.45, p<0.05) with **(A)** the TLR cocktail induced innate immune response (V1) and **(B)** the T cell immune response (V3) **(C)** Representative correlation plots of metabolites from V1 and V3 that correlate with the cytokine release responses from TLR stimulated innate, and peptide pool stimulated T-cell immune responses. Each dot represents an individual sample. Spearman correlation coefficient (R) is indicated. GZB, granzyme B; fc, fold change – obtained by dividing the cytokine concentration in stimulated wells by unstimulated wells;

## DISCUSSION

No other viral disease in the history of humanity has been studied and reported to the extent of COVID-19. And neither has any disease in recent history caused the socio-economic devastation of life on such a global scale^39^. Several integrative studies have attempted to understand the immune correlates of protection against COVID-19^40-42^. These studies have shown that the late induction of T cell responses^43^, impaired type I interferon response^44, 45^, and weakened innate immune responses are immune hallmarks of severe COVID-19^12, 40^.

In this study, the samples were collected during the first wave of COVID-19 (between 07th July and 04th September 2020) before the existence of delta, and omicron VoCs, which allowed us to determine how T cells primed with the ancestral strain react with spike antigen of delta and omicron variants. The magnitude of the antigen-specific T cell response against the ancestral spike was similar to the previous reports^7, 10^. Furthermore, we observed that the T cell response elicited during the active ancestral SARS-CoV-2 infection persists but is relatively reduced against the delta and omicron variants compared to the ancestral spike. The extent of reduction in the magnitude of the T cell response against VoCs is more prominent in our cohort than in previous reports in vaccinated individuals ^4, 5, 46^. Such observation could be because the overall magnitude of T cell response is low in acute samples^7^ and, therefore, may have resulted in a limited scope for cross-reactivity against the VoCs.

Given the importance of T cell responses in limiting disease severity and protection against emerging variants, the next generation of vaccines must primarily focus on the magnitude of T cell responses generated upon vaccination. However, limited studies have investigated the factors contributing to robust T cell immune responses during COVID-19. For the differentiation and activation of functional T cells, the antigen-presenting cells (APCs) must provide three signals to the T cells. One is via direct TCR-MHC interaction, the second is through cognate interaction of adhesion molecules, and the third is by cytokine signaling^11^. Therefore, the APCs play a critical role in shaping the T cell response during exposure to a foreign antigen. An integrative study reports that type I interferon response does not correlate with the antiviral T cell responses^14^. In contrast, another combinatorial study reported a temporary increase of IFN-β and IP10/CXCL10 levels associated with a dominant SARS-CoV-2-specific CD4 T cell response^13^. However, none of these combinatorial reports studied the TLR-specific functional innate immune response, which is more immune cell-specific and functional in nature than the systemic plasma levels^12^.

Increased plasma IL-6, TNF-a, and IL-1β are characteristic of severe COVID-19 patients ^47^. However, it has now been well established that the source of these proinflammatory cytokines is not the innate immune cells but the non-immune host tissues such as the lungs^12^. Furthermore, in severe COVID-19, the innate cell expression of these proinflammatory cytokines is instead significantly reduced^12^. Our investigation of innate and T cell response across the early stages of infection showed that the PBMCs from individuals with a strong proinflammatory innate immune response also exhibit a robust T cell response against SARS-CoV-2 infection. IL-18 is a proinflammatory cytokine that primarily promotes a type 1 response and activates established Th1 cells in IFN-γ production^48^. However, the correlation of proinflammatory Th17 differentiating cytokines, IL-23, and IL-1β ^49^, with the Th1 response in the later stages of the disease needs further investigation. Interestingly, we recently reported that IL-17A transcripts were higher at the early stage (2-4 days post-infection) of SARS-CoV-2 infection in hamsters^50^. Therefore, it may be interesting to investigate further if any Th1/Th17 plasticity^51^ may play a role in the development of T cell response during COVID-19. Such investigations are fundamental for designing adjuvants that trigger a robust and long-lasting memory T cell immunity.

There is an orchestrated symphony between the metabolism and immune responses where both players harmonize^52^. Although other groups have reported the longitudinal metabolomic signature^53^ and single-cell transcriptomic landscape in COVID-19 patients^41, 54, 55^, no study correlated the functional innate and T cell response with the metabolic landscape. Our analysis found a significant correlation between the SARS-CoV-2 specific immune responses and the metabolites involved in pathways such as arginine biosynthesis and arginine and proline metabolism. L-Arginine is an immunomodulatory metabolite that plays a critical role in the pathways involving inflammation, immune regulation, and via its metabolism leading to the generation of nitric oxide (NO), which plays an intrinsic role in Th cell activation, differentiation, survival and proliferation^56-58^. Notably, metabolites involved in arginine biosynthesis have been reported to be significantly altered in severe COVID-19 patients^53^ and their levels are inversely correlated to the severity of COVID-19^59^. Shen et al. reported that the metabolites involved in arginine metabolism were significantly reduced in the COVID-19 patients; in contrast, we observed that these metabolites were elevated in COVID-19-infected individuals compared to the healthy controls (**Fig. S5**). Such differences may be attributed to our study’s predominantly asymptomatic or mildly symptomatic cohort, which further underlines the significance of metabolites involved in arginine metabolism in COVID-19 severity. Furthermore, a randomized, double-blind, placebo-controlled trial on patients hospitalized for severe COVID-19 reported that adding oral l-Arginine to standard therapy in patients with severe COVID-19 significantly decreases the length of hospitalization at ten days after starting treatment^60^. These reports appear to go against the previously suggested strategy of l-Arginine depletion in COVID-19, based on the premise that l-Arginine is a key nutrient essential in the life-cycle of SARS-CoV-2^61^. We speculate that given the role of Arginine biosynthesis metabolites in inducing a more potent T cell response, l-Arginine supplementation could also be beneficial in COVID-19 vaccination and booster doses to induce long-term immunity.

The immunomodulatory effects of metabolites such as Adenosine^62^, Itaconate^63-65^, and L-Proline^66^ have been reported independently or in the context of disease pathogenesis. Enrichment of these metabolites as correlative markers of anti-SARS-CoV-2 immune responses further underlines their importance in immunomodulatory mechanisms.

To conclude, our study highlights the importance of the T cell response against the emerging variants of SARS-CoV-2. We show how PBMC-derived innate responses are critical for generating a robust T cell response and how plasma metabolies correlate with the innate and asaptive immune responses. How these metabolites could act as potential immunomodulator needs further examination.

## Acknowledgment

We acknowledge the funding through the Mission COVID Suraksha grant (BT/CS0010/CS/02/20) – from BIRAC, Department of Biotechnology, for enabling this study. We thank all the volunteers who donated their blood samples for this study. We sincerely thank the staff and students of the Translational Health Science and Technology Institute (THSTI; Faridabad, India) for helping to ensure the smooth conduct of the study.

## Availability of Data and Material

The datasets generated during and/or analyzed during the current study are available from the corresponding author, AA, on reasonable request.

## Authors’ contributions

Conceptualization, Akshay Binayke, Deepak Rathore and Amit Awasthi; Data curation, Shailendra Mani, Yashwant Kumar, Deepak Rathore and Amit Awasthi; Funding acquisition, Deepak Rathore and Amit Awasthi; Investigation, Shailendra Mani, Tripti Shrivastava, Yashwant Kumar, Deepak Rathore and Amit Awasthi; Methodology, Akshay Binayke, Aymaan Zaheer, Jyotsna Dandotiya, Sonu Kumar Gupta, Shailendra Mani, Manas Tripathy, Upasna Madan, Tripti Shrivastava and Yashwant Kumar; Project administration, Anil Pandey, Deepak Rathore and Amit Awasthi; Supervision, Anil Pandey, Deepak Rathore and Amit Awasthi; Visualization, Akshay Binayke; Writing – original draft, Akshay Binayke and Aymaan Zaheer; Writing – review & editing, Amit Awasthi.

## Ethics Approval

All the experiments were performed according to the suggested guidelines of the Institutional Ethics Committee (Human Research) of THSTI and ESIC Hospital, Faridabad (Letter Ref No: THS 1.8.1 (97) dated 07th July 2020). Human blood samples were collected from COVID-19 patients and healthy individuals after the written informed consent. Individuals were enrolled in this study based on the inclusion/exclusion criteria prescribed by the Institutional Ethics Committee (Human Research) of THSTI.

## Consent to Participate

Informed consent was obtained from all individual participants included in the study.

## Consent to Publish

Not Applicable.

