## Supplementary Material for "Proinflammatory innate cytokines and metabolomic signatures shape the T cell response in active COVID-19"

**MATERIALS AND METHODS**

**Expression and purification of RBD protein of wildtype SARS CoV-2 and Variants:**

RBD protein of ancestral and VoCs SARS-CoV-2 RBD proteins were expressed and purified from Expi293 cells (Thermo Fisher) by performing site-directed mutagenesis one by one for N501Y (B·1·1·7), K417N, E484K and N501Y (B·1·351), and E484Q, L452R (B·1·617.2) as described previously (*1*). The culture supernatant of transfected cells with secreted protein was harvested after six days of transfection by centrifugation of the culture supernatant and loaded onto Ni-NTA agarose (Qiagen) for affinity purification. The purified fractions were then pooled, concentrated, and further purified through size exclusion chromatography employing a Superdex 200 Increase 10/300 GL column (GE Healthcare), equilibrated in PBS. The purified protein was stored at −80 °C until further use.

**Peripheral Blood Mononuclear Cells (PBMC) Isolation:**

Venous blood from healthy participants and COVID-19 patients were collected in CPT^TM^ tubes (BD Biosciences) and centrifuged at 1500g for 25min at 25˚C within 2h of drawing of blood draw. Plasma was separated from the upper layer, aliquoted, and stored at -80˚C until further use. The layer of PBMCs was transferred into a 15ml tube and washed with sterile ice-cold PBS. The total PBMCs were counted on a hemocytometer after staining with Trypan Blue and were resuspended in freezing media (90% FBS + 10% DMSO) followed by storage at -80°C for at least 24h, before being transferred to liquid nitrogen.

**Identification of mutations in dominant class I and class II epitopes of SARS-CoV-2 spike:**

We curated a list of dominant epitopes on the basis of previous published literature and the epitopes listed on IEDB database (*3*). The dominant epitopes were defined as previously described in (*4*). Only those dominant epitopes that are mutated in delta or omicron were further shortlisted and summarized in **Table S2.**

**THSTI In-house RBD IgG and IgM ELISA:**

ELISAs to detect IgG and IgM binding to RBDs were performed as previously described (*2*) with some modifications. In-house RBDs of the Ancestral (Wuhan Isolate), Alpha (B.1.1.7), Beta (B.1.135), and Delta (B.1.617.2) variant S proteins were used at 2 µg/ml. A BioTek 405 LS microplate washer carried out all washes. 250μl of blocking solution was used, and the incubation for the blocking step was carried out for 2 hours at RT, after which the plate was washed once with PBS with 0.1% Tween 20 (PBST). All other incubations were carried out at RT for 30 min. After the addition of plasma samples, all washes were performed five times. The plasma samples were diluted with 3% skim milk at 1:50 dilution for IgG and 1;100 for IgM. Tracer antibody IgG (anti-human IgG, Jackson Immunoresearch Laboratory, USA) and IgM (goat anti-human IgM, INVITOGEN) were diluted at 1:10000 in PBST with 3% skim milk. For detection, 100ul of TMB substrate was added to each well followed by the addition of 100μl of stop solution (1N H_2_SO_4_). Absorbance was measured on a BioTek Synergy HT Microplate reader.

A positive convalescent and negative control sample were added to each plate at a single point concentration for normalization. The assay results were normalized by dividing the blank subtracted readings of each sample by negative control to obtain the fold change reading. Cut-off values were calculated with the following formula: Cut-off = Average fold change value of negative samples (pre-pandemic plasma samples) + 3* SD of fold change value of negative samples.

**Virus Neutralization Assay:**

Titres for the virus microneutralization assay were estimated as previously detailed (*1*). In short, we seeded Vero E6 cells in a 96-well plate, at 30,000 cells/ well. 75 μl serum samples, after heat inactivation, were serially diluted in growth medium containing 2% heat-inactivated FBS, from 1:20 to 1:2560. 75 μl SARS-CoV-2 (with pre-determined dilution for producing 50-150 microplaques) was added, and the solution was kept in a 5% CO2 incubator for 1 h at 37oC. Virus-serum mixtures were subsequently added to Vero E6 cells before another incubation for virus adsorption at 37°C for 1 h. The viral inoculum was then removed before 1·5% carboxymethylcellulose (Sigma-Aldrich) in growth medium containing 2% heat-inactivated FBS was used to overlay the cells, which were then incubated at 37°C in a 5% CO 2 incubator. The cells were fixed with a formaldehyde solution 24 hours post infection, stained for 1 hour with 1:4000 diluted anti-spike RBD antibody (Sino Biologicals), then for 1 hour by 1:4000 diluted HRP-conjugated anti-rabbit antibody (Invitrogen). The cells were washed with PBS, incubated for 10 minutes with TrueBlue substrate (Sera Care), then washed with sterile MilliQ water. After the plates were air-dried, the microplaques were quantified on an AIDiSPOT reader (AID GmbH, Strassberg, Germany), using AID EliSpot 8·0 software. Calculation of 50% neutralization values was done by four-parameter logistic regression on GraphPad Prism 7·0e software. Experiments with SARS-CoV-2 were solely done in biosafety level 3 labs:

| **No.** | **Virus Name** | **Virus Details** | **Accession No.** |
| --- | --- | --- | --- |
| 1 | SARS-CoV-2 Wuhan Isolate | Isolate USA-WA1/2020, NR-52281 | GenBank: MN985325, GISAID:  EPI_ISL_404895, GenBank:  MT020880 |
| 2. | SARS-CoV-2 (delta) | THSTI_287 | GenBank: MZ356566.1 |

**Flow Cytometry:**

For flow cytometry, the stimulated PBMCs were stained for viability and surface markers (CD4 FITC, 1:100; CD8 BV510, 1:100; CD69 PE-CF594 1:100; CD137 BV-605 1:100; OX-40 PE-Cy7 1:100, all BD Biosciences) in FACS buffer (PBS supplemented with 2% FBS (Gibco)) for 40 min at 4 °C. Next, cells were fixed and permeabilized using the Cytofix/Cytoperm kit (BD Biosciences) according to the manufacturer's instructions. Intracellular staining was performed by incubating the cells in Perm/Wash buffer supplemented with antibodies for 30 min at 4 °C (IFNγ PE, 1:50; IL-2 BV711, 1:50; TNFα BB700, 1:50; GranzymeB (GZB) PE 1:50; Perforin APC 1:100 all BD Biosciences). Samples were acquired on a fluorescence-activated cell sorter (FACS) Symphony^TM^ instrument (BD Biosciences) using BD FACSuite software version 1.0.6 and analyzed with FlowJo software version VX (FlowJo LLC, BD Biosciences).

**Metabolomics analysis LC-MS/MS reverse phase:**

100% methanol was added to 100 µl of serum samples to extract metabolites. After being vortexed for 1 min, the resultant mixture was kept on ice before protein precipitation. The sample underwent centrifugation at 10000 rpm for 10 min at 4ºC, before the supernatant was collected and dried for 20-25 min by speed vacuum. The samples were dissolved in 15% methanol in water (v/v) and then stored at -80ºC till they were analyzed.

**Measurement of metabolites.**

The sample metabolite data acquisition was performed using an Orbitrap Fusion mass spectrometer (Thermo Scientific) coupled with the heated electrospray ion source. Data acquisition was performed as described previously (*5, 6*) with minor modifications described below. The mass resolution for MS1 acquisition was set at 120,000, whereas for MS2 mode, the mass resolution was 30,000. The mass range for data acquisition was 60–900 da. The extracted compounds were separated on a UPLC ultimate 3000 and the data was acquired on HSS T3 and HILIC columns. The Solvent A was distilled water and B was methyl alcohol with both solvents supplemented with 0.1% formic acid. The elution gradient range from 1% B to 95% B over 10 min with a flow rate of 0.3 ml/min. The sample injection volume was set at 5ul. A pooled quality control (QC) sample was run post every five samples to keep track of signal variation and drift in mass error.

**Data Processing for Metabolite Measurement**

The LC/MS obtained data were processed using the Progenesis QI for metabolomics (Nonlinear dynamics, a Waters Company) software using the default setting. The untargeted workflow of Progenesis QI was employed to execute the retention time alignment, feature detection, deconvolution, and elemental composition prediction. The Metascope plug of Progenesis QI was used for the in-house library with retention time, accurate mass, and fragmentation pattern for database search. We also used the spectral library available online for further identity confirmation. Peaks with a coefficient of variation (CV) of less than 30% in pool QC samples were kept for further data analysis. The cut-off for retention time match was 0.5 min, and the spectral similarity was more than 30% fragmentation match in Progenesis QI. Additionally, each detected feature was manually verified to select the appropriate peaks.

**Table S1: Donor Characteristics**

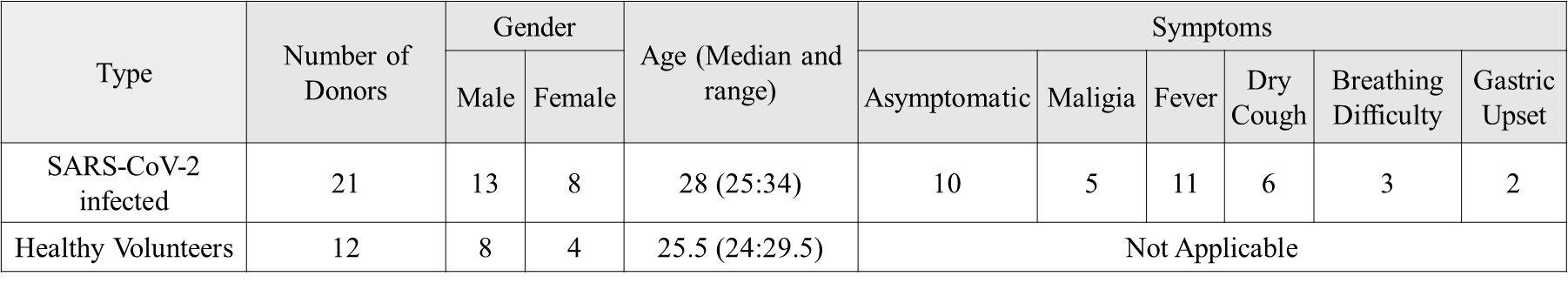

**Fig. S1: Gating Strategy**

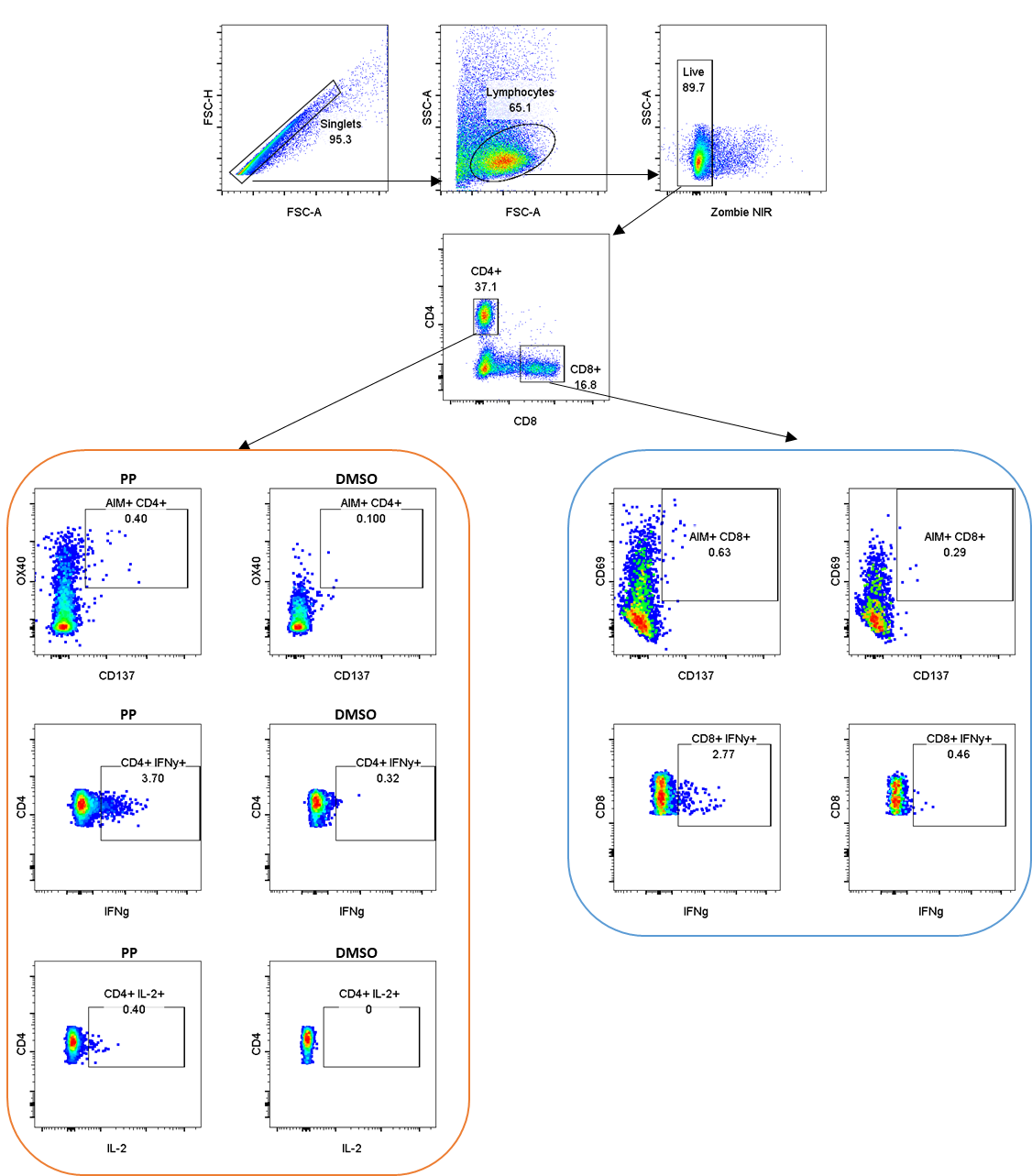

**Fig S2: SARS-CoV-2 virus-specific innate immune responses** determined by stimulating the PBMCs from early stages of COVID-19 positivity (V1: day 0-3) with TLR agonists R848 and poly I:C for 24h. The cytokine released in the culture supernatant was calculated by Luminex’s bead-based cytokine release assay. Each dot represents an individual sample and the paired samples were analysed using the Wilcoxon Signed-Rank test. Bars represent mean values for each time point.

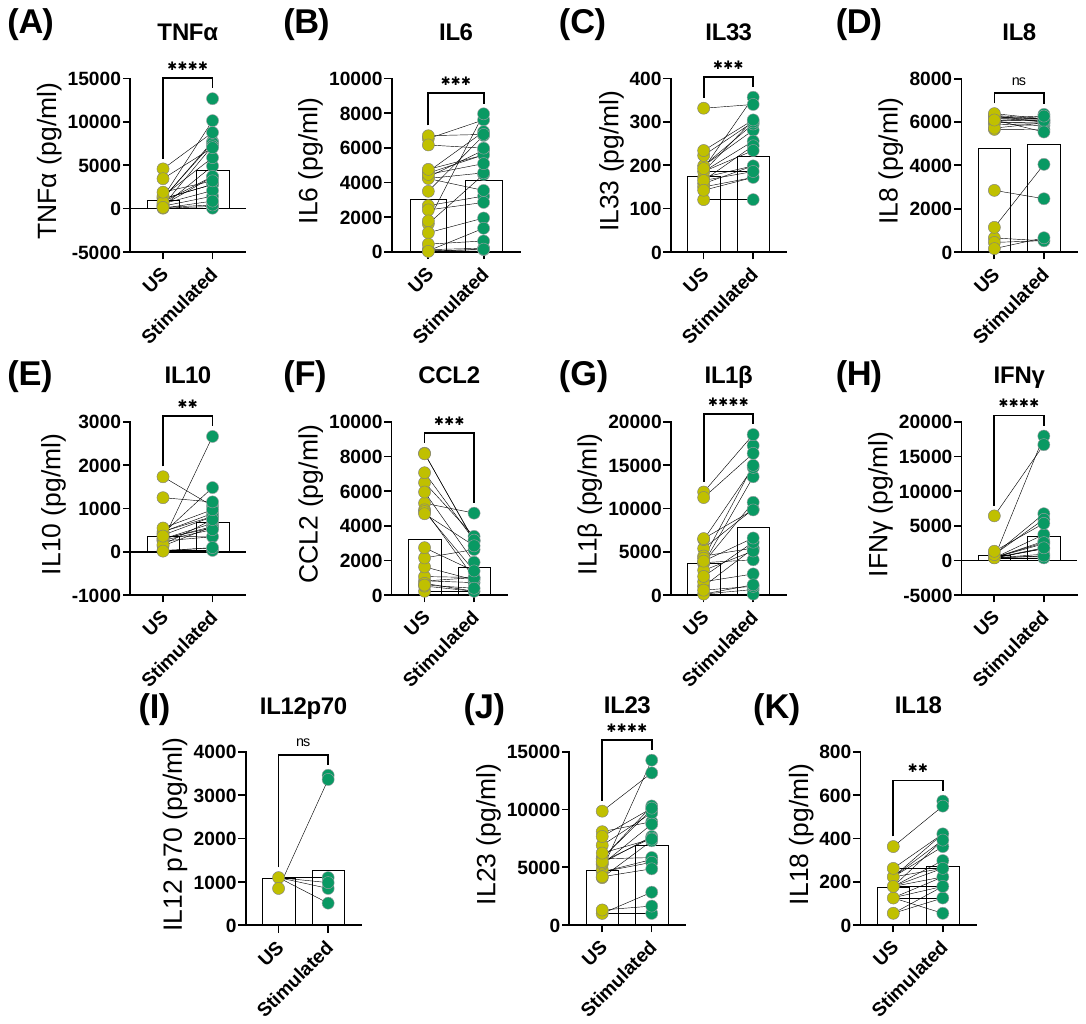

**Fig. S3: Longitudinal Analysis of IgG and IgM Response against the SARS-CoV-2 during acute COVID-19 Infection.** The longitudinal anti-RBD IgG responses were evaluated by performing ELISA against the RBD proteins of **(A)** Delta (B.1.617.2) **(B)** Beta (B.1.315) **(C)** Alpha (B.1.1.7) variants of SARS-CoV-2. The anti-RBD IgM titers were also calculated against the RBD proteins of **(D)** WT (Ancestral - Wuhan isolate) **(E)** Delta (B.1.617.2) **(F)** Beta (B.1.315) **(G)** Alpha (B.1.1.7). All data, represented as ratio-converted ELISA reads to a pool of pre-pandemic negative control samples (relative ratio), were plotted using violin plots. The dots represent the relative ratios of each individual sample. Tukey’s multiple comparison test was employed.  n.s = not significant, *= p ≤ 0.05, ** = p ≤0.01.

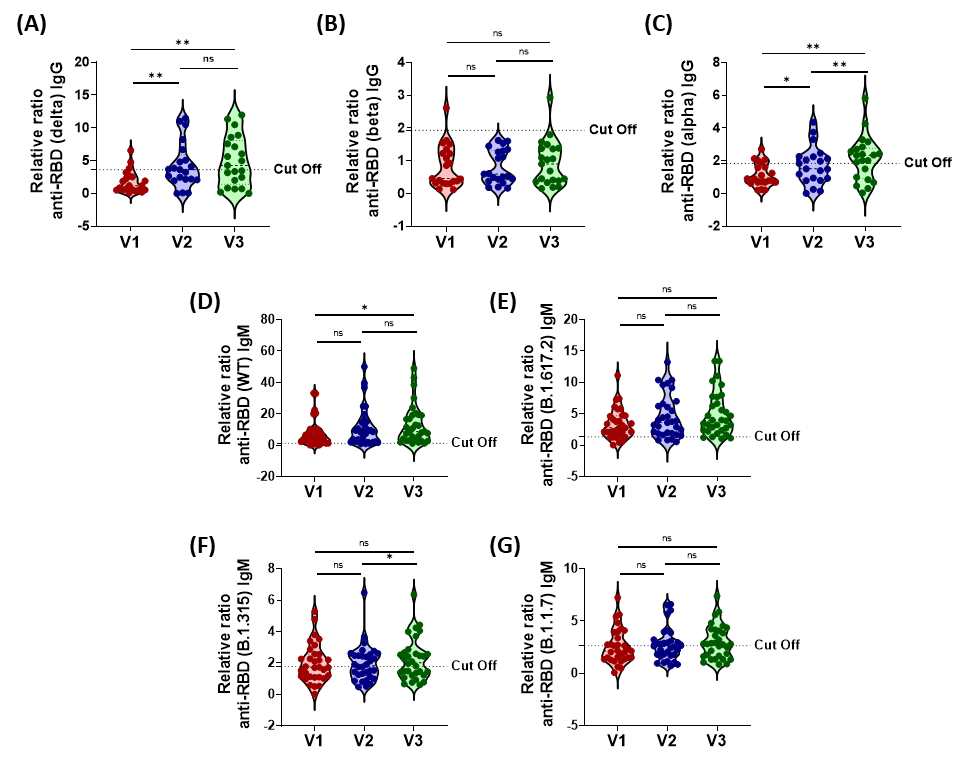

**Table S2: List of dominant epitopes** shortlisted from the literature and IEDB database that are mutated in Omicron and Delta variants. The mutations are marked in red colour. We found that 96% each of the dominant CD4 and CD8 epitopes are conserved against the delta spike, whereas 80% and 82% of CD4 and CD8 dominant epitopes, respectively, are conserved against the Omicron spike

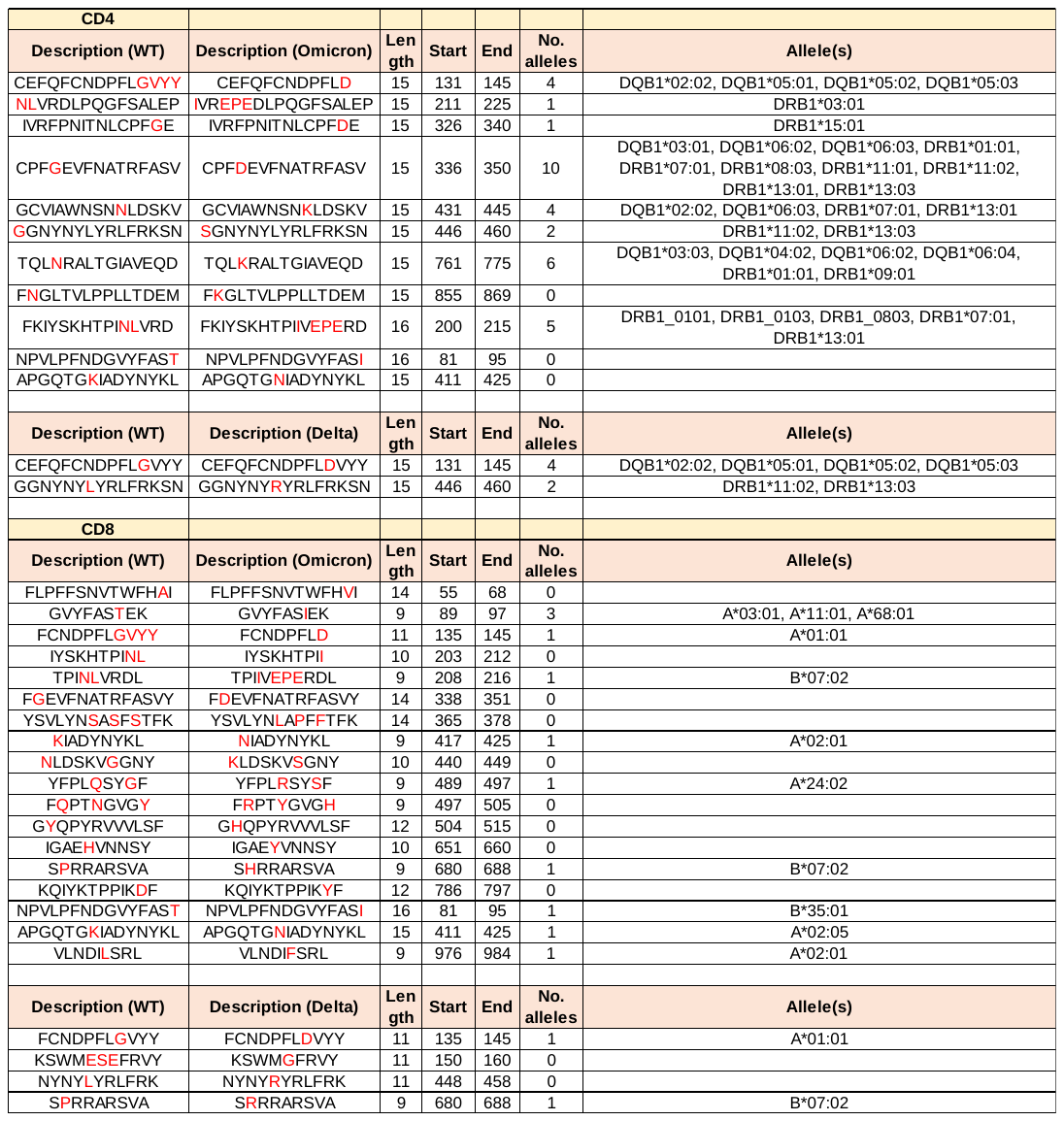

**Fig. S4:** Normalized Box plots and kernel density plots using the online tool Metaboanalyst 5.0. The density plots are based on all the samples uploaded. Strategy for normalization: Data transformation: Log10 Normalization; Row-wise normalization: Normalization to constant sum; Data scaling: Pareto Scaling.

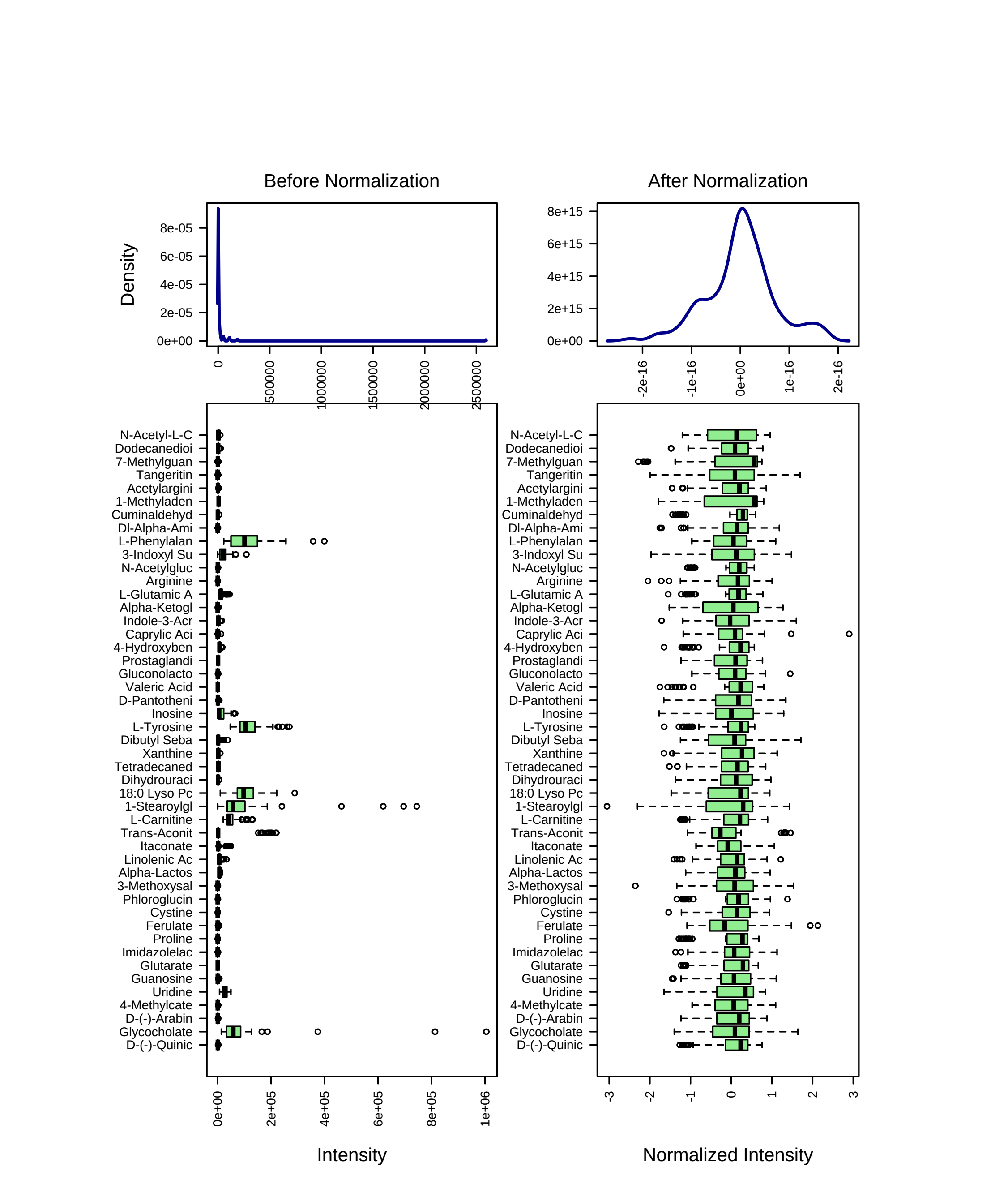

**Fig. S5: Normalized Box plots of representative plasma metabolites** in healthy volunteers (control) and at early stage (V1) and late stage (V3) of COVID-19 disease. Each dot represents an individual sample. The bold black line represents the median values, box represents the inter quartile range and the vertical line represents the range of normalized values of the peak intensities of the metabolites estimated by mass spectrometry.

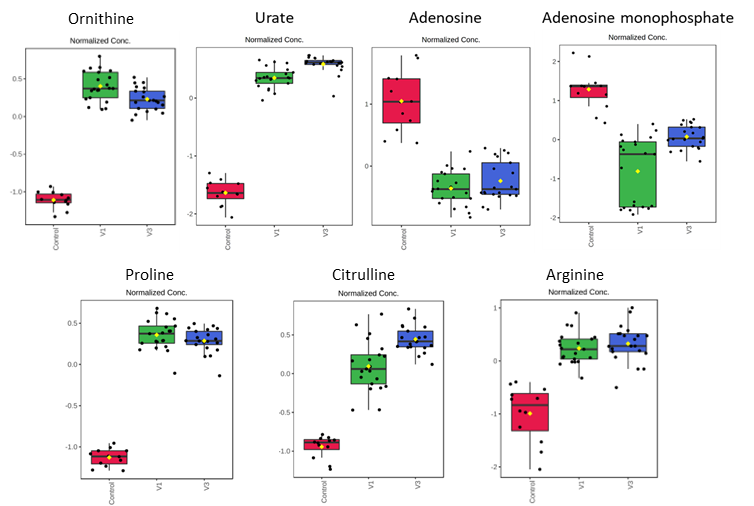

**Fig. S6: SARS-CoV-2 Spike Peptide Pool Specific Cytokine Release by PBMC samples of V3.** PBMCs from late stages of COVID-19 positivity (V3: day 14) with peptide pools for 24h. The cytokine released in the culture supernatant was calculated by Luminex’s bead-based cytokine release assay. Each dot represents an individual sample and the paired samples were analysed using the Wilcoxon Signed-Rank test. ****** p<0.01; * p<0.05; ns, not significant.

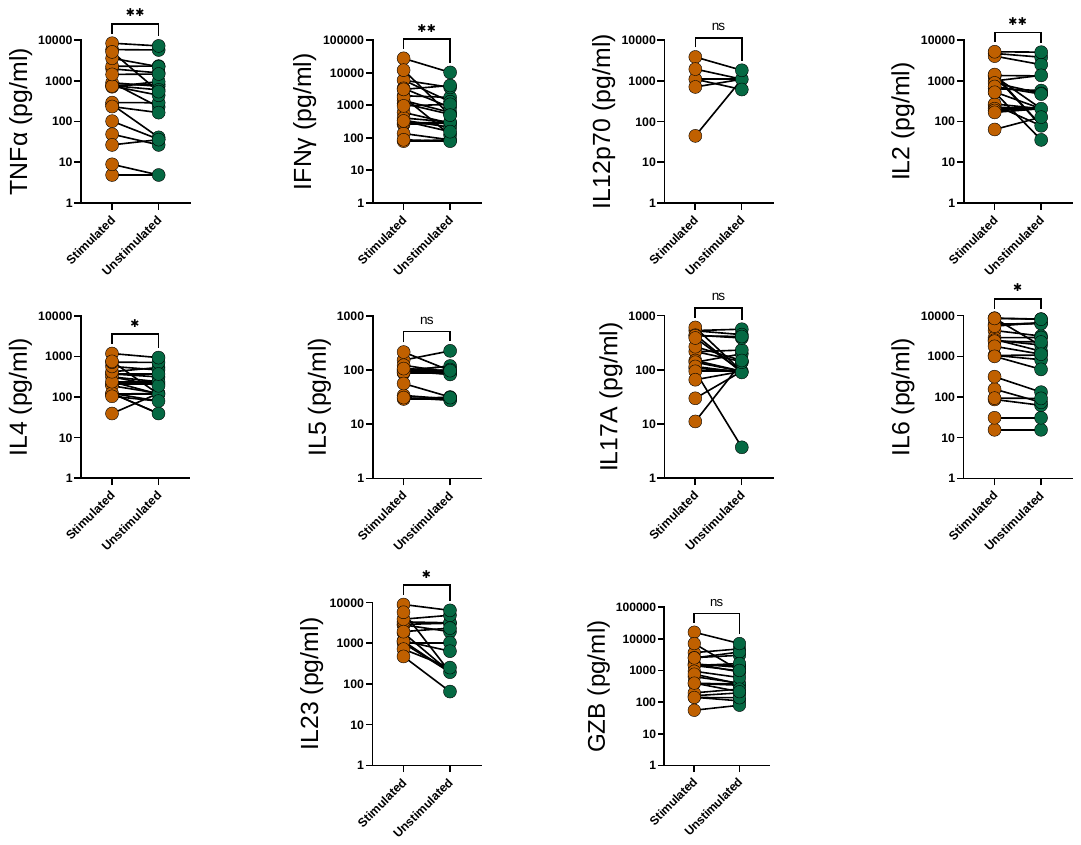

**Fig. S7. Longitudinal dynamics of antigen-specific cytokine release by PBMCs during COVID-19 infection.** Graphs represent longitudinal cytokine released by paired PBMC samples from the same subject upon stimulation with SARS-CoV-2 spike peptide pool for 18-20h. Each dot represents an individual sample, and the paired samples were analyzed using the two-sided Wilcoxon Signed Rank test. Bars represent mean values.

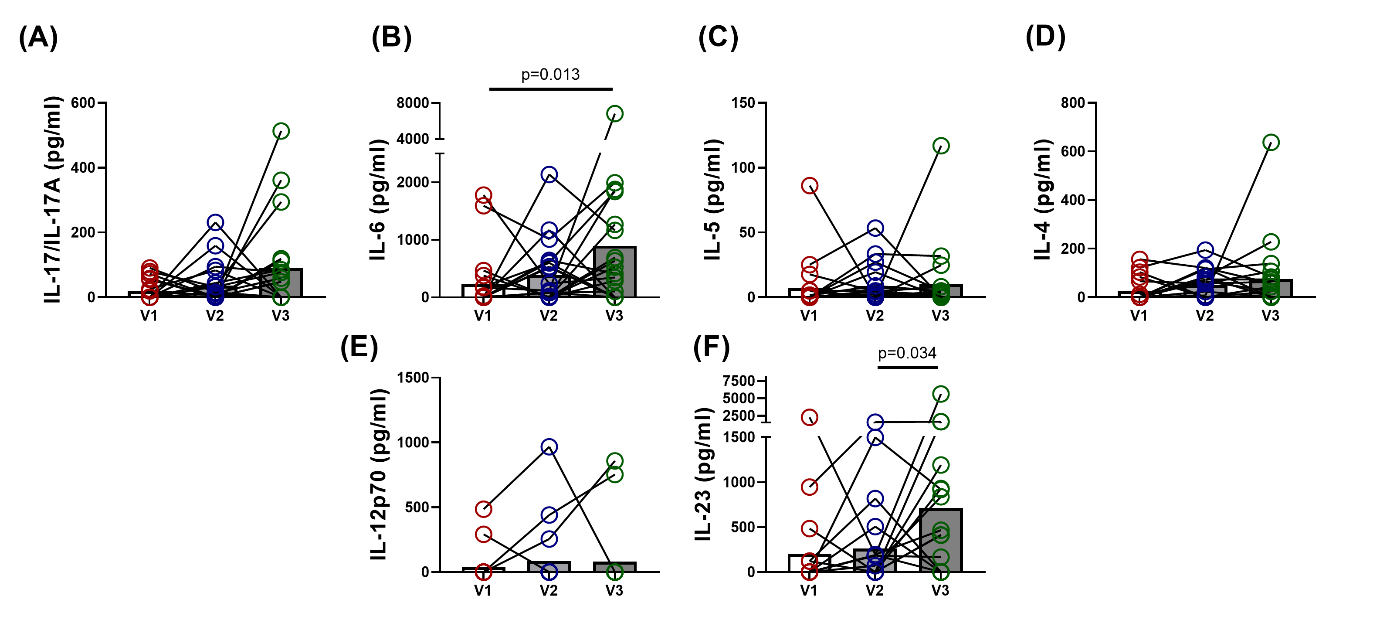
